# An omnidirectional visualization model of personalized gene regulatory networks

**DOI:** 10.1101/644070

**Authors:** Chixiang Chen, Libo Jiang, Ming Wang, Yaqun Wang, Biyi Shen, Zhenqiu Liu, Zuoheng Wang, Wei Hou, Scott A. Berceli, Rongling Wu

## Abstract

Gene regulatory networks (GRNs) have been widely used as a fundamental tool to reveal the genomic mechanisms that underlie the organism’s response to environmental and developmental cues. Standard approaches infer GRNs as holistic graphs of gene co-expression, but such graphs cannot quantify how gene-gene interactions differentiate among organisms and how they alter structurally across spatiotemporal gradients. Here, we develop a generalized framework for inferring informative, dynamic, omnidirectional, and personalized GRNs (idopGRNs) from routine transcriptional experiments. This framework is constructed by a system of quasi-dynamic ordinary differential equations (qdODEs) derived from the combination of ecological and evolutionary theories. We reconstruct idopGRNs from a clinical genomic study and illustrate how network structure and organization affect surgical response to infrainguinal vein bypass grafting and the outcome of grafting. idopGNRs may shed light on genotype-phenotype relationships and provide valuable information for personalized medicine.

## Introduction

Gene regulatory networks (GRNs) have been thought to operate as the genomic mechanisms that guide the organism’s response to changes in their environment^1,2^. One promising subject of research in modern biology and translational medicine is how to infer biologically realistic and statistically robust GRNs from increasingly available transcriptional data and link them to physiological, pathological, and clinical characteristics^3-5^. A number of statistical approaches, such as Boolean networks^6^, Bayesian networks^7^, mutual information theory^8,9^, and graphical models^10^, have been developed for network inference, and these approaches visualize GRNs as probabilistic, undirected or unidirectional graphs, where each node represents a gene and edges depict relationships between genes. However, such graphs may not be sufficiently informative for charting the topological structure of a GRN because genes may regulate and also be regulated by other genes, with regulations in various signs and strengths and varying across time and space scales^3,11^.

As the time generalization of Bayesian networks, dynamic Bayesian networks (DBNs) can code cyclic, causally directed, and probabilistic interactions into networks through temporal interdependence, but they are often puzzled by the choice of granularity when time spaces vary^12-14^. When gene networks are modeled by a system of time-derivative ordinary differential equations (ODEs), all these issues can be mostly addressed^15-18^. The successful use of such ODE-based networks is, however, impaired by two factors: (1) parametric dynamic modeling, which is difficult to justify, given that gene expression is often stochastically fluctuated^19,20^ and alters across discrete regimes, such as cell/tissue types and medical treatments^21^, and (2) the requirement of high-density temporal expression data over a time course^22^. Gene networks are regarded as temporal or spatial snapshots of biological processes^23^, but no existing approaches can contextualize how GRNs change structurally and functionally in response to developmental and environmental cues. More importantly, most approaches can only identify an overall network from a set of cross-sectional or longitudinal data, largely limiting the use of GRNs as a personalized tool for clinical diagnosis and prediction of individual subjects in the era of precision medicine.

Here, we develop a statistical framework for inferring informative, dynamic, omnidirectional, and personalized GRNs (idopGRNs) from standard genomic experiments. An ***informative*** network should encapsulate bidirectional, signed, and weighted edges that facilitate the interpretation and interrogation of gene-gene interactions. A ***dynamic*** network can monitor how the pattern of gene co-expression alters in response to environmental and developmental change. An ***omnidirectional*** network codes all possible gene interactions but ensuring its sparsity and stability. Because of different genetic backgrounds, specific individuals may develop and use their ***personalized*** networks to regulate any phenotypic change. To recover such idopGRNs, we integrate elements of distinct disciplines into a unified framework by which expression data from multiple individuals under distinct treatments, monitored at several key time points and/or across spaces, can be assembled, modeled, and analyzed. We virtualize idopGRNs as an ecological community composed of many species, in which the expression level of each gene, corresponding to the abundance of each species, is determined by its niche and niche differences collectively stabilize the whole network through gene-gene interactions in a way similar to interspecies interactions^24-26^. We integrate the niche theory of biodiversity and evolutionary game theory to derive a system of quasi-dynamic ordinary differential equations that model gene networks across individuals. The implementation of variable selection helps to define and select a subset of the most significant genes that regulate a focal gene, which enables the inference of sparse but omnidirectional networks. To test and validate our approach, we analyzed genomic data of circulating monocytes from human infrainguinal vein bypass grafting, aimed at treating lower extremity arterial occlusive disease^27^, and reconstructed graft- and outcome-perturbed idopGRNs. The usefulness of our approach is further validated by a second vein graft experiment for rabbits^28^. In both cases, quantitative comparison of GRN structure and organization between different outcomes and across times provides a mechanistic understanding of vein bypass graft success vs. failure.

### Theory Construct

The theory for reconstructing idopGRNs is interdisciplinary, founded on the seamless integration of community ecology, evolutionary biology, and network science through mathematical and statistical reasoning. Each discipline contributes its distinct elements to a unified framework of statistical inference for gene networks.

### Niche theory of biodiversity

The concept of niche was first defined by Elton^29^ to describe the ecological components of a habitat related to a species’ tolerance and requirement. This concept has been generalized to explain biodiversity and species coexistence patterns in ecological communities^30^. A gene network, residing in any biological entity, such as a cell, a tissue, or even an individual, can be viewed as an ecological community, in which the expression level of a constituent gene corresponds to the niche occupied by a species and niche differences form community diversity and stability. From a community ecology perspective, the total expression amount of all genes in the network reflects the carrying capacity of the entity to sustain indefinitely these genes and supply them with essential resources or energy for their function^31^, which are a mixture of many unknown factors. We define the total expression level of all genes on an entity as the expression index (EI) of this entity. This concept, similar to environmental index coined to describe the overall quality of site in terms of the accumulative growth of all plants^32,33^, can describe the overall occupation of all genes to the entity. By aligning EI values in an ascending order, we can convert discrete entities to a series of continuous variables that help establish a system of ordinary differential equations (ODEs).

In an ecological habitat, each organism needs to respond to the distribution of resources and competitors and it in turn alters those same factors^34^. For example, an organism would grow fast when resources are abundant, or when predators or parasites are scarce, and may limit access to resources by other organisms or provide a food source for predators. The types and numbers of environmental variables constituting the dimensions of a habitat vary from one species to another and the relative importance of particular environmental variables for a species may vary according to the geographic and biotic contexts^35^. Thus, based on the niche theory of biodiversity, the relationship of the abundance of a particular species (part) with the total abundance of all species (whole) across graded habitats can potentially describe and predict the inherent compositional structure of an ecological community and its response to environmental change. This part-whole relationship, governed by the power scaling theory, has been observed to pervade biology; For example, the power equation can well explain how total leaf biomass scales allometrically with whole-plant biomass across different plants^36,37^ and how brain size of animals scales with whole-body mass across animals^38,39^. We introduce this power scaling theory to model how the expression of individual genes (part) scales with the total expression of all genes across EIs through a system of ODEs.

### Evolutionary game theory of gene expression

In an ecological community where many species coexist, a species may adopt a cooperative or competitive decision to maximize its chance to access to resources^40^. This phenomenon has also been well recognized at the cell level in both humans and rats^41,42^, by which a cell determines a goal-directed decision-making based on its accrued knowledge of the environment. In an elegant study of stress impact, Friedman et al.^43^ identified the cells and networks that enable a rodent to choose an appropriate strategy of responsiveness after evaluating possible costs and benefits. Such rational choice reasoning may also guide how genes, located in the same cell, promote or inhibit each other in a complex network. In other words, gene-gene interactions can be modeled as a game in which one player may choose to compete or cooperate with its opponents in a quest to maximize its payoff. Classic game theory, pioneered by mathematical economists^44^, suggests that such choices are not arbitrary, but rather include a rational judgement based on a gene’s own strategy and the strategies of other genes. However, it is extremely difficult or impossible to interrogate the rationality of genes, making a direct application of classic game theory to gene network inference infeasible. To address this issue, we introduce evolutionary game theory, a combination theory of game theory and evolutionary biology^45^, which does not rely on the rationality assumption when it is used to study community dynamics and evolution. In an evolving population, any strategy used by an individual to maximize its payoff would be constrained by strategies of other individuals that also strive to maximize their own payoffs and, ultimately, this process through natural selection would optimize the structure and organization of the population, making it reach maximum (best response) payoff^45^.

### Mathematical integration of evolutionary game theory and niche biodiversity theory

Suppose we initiate a standard genomic experiment (Fig. 1**A**) involving *S* treatments, each with *n*_*s*_ (*s* = 1, …, *S*) subjects, measured for *m* genes and *p* phenotypic traits at a series of time points (*t*_0_, *t*_1_, …, *t*_*T*_), where *t*_0_ denotes pre-treatment and *t*_1_, …, *t*_*T*_ denote post-treatment. We call a subject from a treatment measured at a time point a “sample.” Thus, we have a total of *N =* (*T* + 1)*n* samples, where 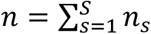 is the total number of subjects from all treatments. Let *M*_*ji*_ denote the expression level of gene *j* (*j* = 1, …, *m*) on sample *i* (*i* = 1, …, *N*). The EI of sample *i* is defined as _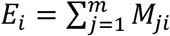_. We line up the *N* samples in the ascending order of EI, which allows us to construct a system of ODEs, expressed as

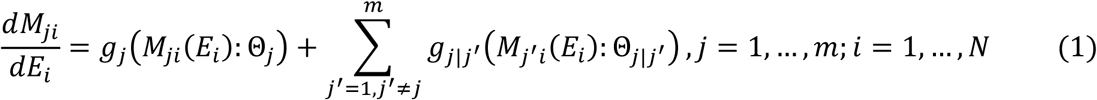

where the change rate of the expression of gene *j* per *E*_*i*_, *M*_*ji*_(*E*_*i*_), at a given sample *i*, is decomposed into the independent expression component, *g*_*j*_(·), specified by unknown parameters *Θ*_*j*_, and the dependent expression component, *g*_*j|j*′_(·), specified by unknown parameters *Θ*_*j|j*′_. The independent component of gene *j* occurs if this gene is assumed to be expressed in an isolated environment, and it is determined by this gene’s intrinsic property. The dependent component of gene *j* is the aggregated effect of all possible other genes *j*′ (*j*′ = 1, …, *m*; *j*′ ≠ *j*) on this gene. General speaking, the independent expression of a gene is determined by its **endogenous** encoding capacity, whereas its dependent expression is under the **exogenous** control. The structure of ODEs in Eq 1 is similar to the generalized Lotka-Volterra equations^46^ with the community matrix replaced by the functions *g*_*j/j*′_ (·) and the time derivative replaced by the EI derivative. Since they are not time based, such ODEs are called quasi-dynamic ODEs (qdODEs). It is straightforward to derive example equations of this type from the multi-gene replicator dynamics. Identifying these functions is a primary focus of research with a secondary effort being in interpretation and analysis of the resulting dynamical system.

**Figure 1.**
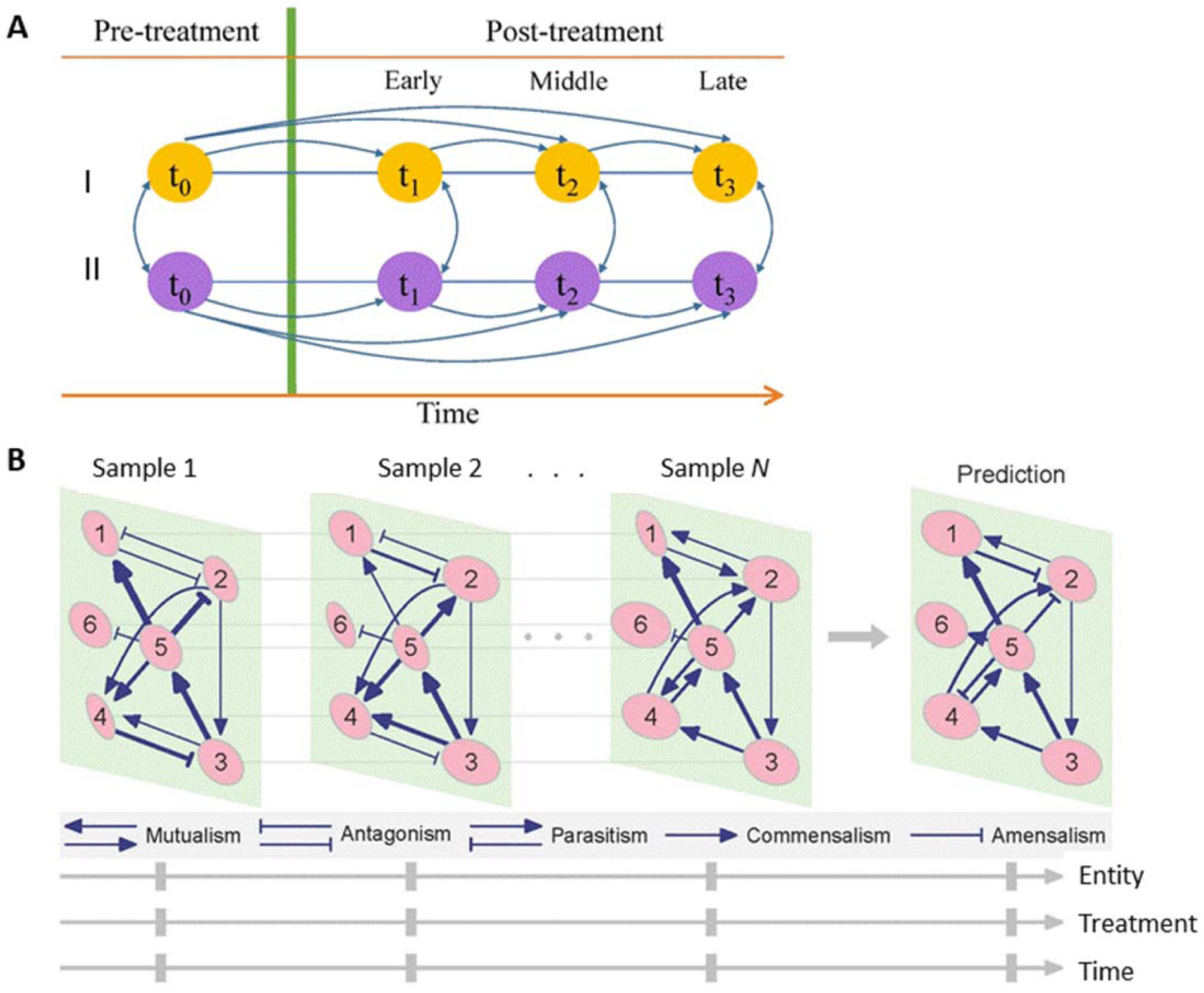
(**A**) Diagram of a standard genomic study including multiple entities under two levels of treatment, I and II. Transcriptomic profiles are monitored at key time points including one before treatment (*t*_0_) and several others at early (*t*_1_), middle (*t*_2_), and late stages (*t*_3_) of response after treatment. (**B**) Illustration of informative, dynamic, omnidirectional, and personalized gene regulatory networks (idopGRNs) among six hypothetical genes from the standard genomic study. idopGRNs vary structurally among samples. For example, genes 1 and 2 are slightly antagonistic in sample 1, moderately antagonistic in sample 2, mutualistic in sample N, and parasitic/altruistic in a predicted sample. The commensalism of gene 5 to gene 1 is strong in samples 1 and N, but weak in sample 2. Because outgoing links are more than incoming links, gene 5 is a social gene in all samples, but the degree of its sociality is different across samples.

### Inferring gene networks

In practice, the number of genes for network reconstruction is commonly very large (e.g., 10^3^– 10^4^), thus if the expression of each gene involves the effects of all other genes, ODEs in Eq 1 will quickly become intractable. Indeed, it is unlikely that each gene performs an interaction with every other gene in the network. By regressing the expression of each gene *j* on the expression of all other genes *j*′ (*j*′ = 1, …, *m*; *j*′ ≠ *j*), we formulate a multiple regression model across samples for variable selection. We implement adaptive LASSO to detect a small set of the most significant genes that affect a focal gene *j* (incoming links), but posing no constraint on the number of genes affected by the focal gene (outgoing links). This procedure enables the reconstruction of a high-dimensional but sparse and stable GRN under the convex optimization formulation (see Online Methods). These GRNs are regarded as **idopGRNs** (Fig. 1**B**) because of their following five major features:

#### (i) Bidirectional, signed, and weighted

Let *G*_*j*_(·) and *G*_*j/j*′_ (·) denote integrals of *g*_*j*_(·) and *g*_*j/j*′_(·) that constitute the system of qdODEs in Eq 1, respectively. Note that, for a focal gene *j*, the number of its incoming links is *d*_*j*_ (<< *m*) after variable selection. The estimate of *G*_*j/j*′_ (·) can help judge in which way gene *j*′ affects gene *j*. If it is positive, negative, or zero, then this suggests that gene *j*′ promotes, inhibits, or is neutral to, gene *j*, respectively. The value of the estimate can quantify the strength of promotion or inhibition. By comparing *G*_*j/j*′_ (·) and *G*_*j* ′ */j*_(·), we can determine whether these two genes reciprocally trigger impacts on each other. Further, we reconstruct a **bidirectional, signed**, and **weighted** graph as the gene network of the sample by considering all possible gene pairs detected from variable selection. The estimate of *G*_*j*_(·) represents how much amount of expression a given gene *j* may intrinsically release, and its value is proportional to the size of a node in the graph.

#### (ii) Dynamic

The amount of dependent expression *G*_*j/j*′_ (·) is a function of *E*_*i*_, suggesting that the dependent amount of gene *j* affected by gene *j*′ can be estimated at any given EI. Thus, we can reconstruct a series of “dynamic” networks across samples. These networks allow geneticists to test how GRNs alter structurally and functionally in response to environmental and developmental cues. These tests can be made locally, i.e., testing how networks differ between two time points of interest under the same treatment or between different treatments at the same time point.

#### (iii) Omnidirectional but sparse

If the number of genes for network reconstruction is large, we should build a high-dimensional set of ODEs that can specify the whole picture of gene interactions in the network. The implementation of variable selection can detect the most significant links to construct a sparse network but still allows all possible realistically existing links to be encapsulated as a whole that underlie the behavior of gene networks. This dimension reduction procedure will become even more valuable since more and more studies attempt to reconstruct regulatory networks from genomic, proteomic, and metabolomics data. A more fine-grained network inferred from these omics data at different levels or through different pathways can reveal previously hidden contributions of gene interactions to cellular processes.

#### (iv) Personalized

The most noticeable advantage of our approach is the ability to pack steady-state expression data into highly informative networks that can currently be inferred only from high-density temporal data. As a function of *E*_*i*_, the independent and dependent expression values of genes can be calculated for any sample from *G*_*j*_(·) and *G*_*j|j*′_(·), respectively. These values enable the inference of sample-specific networks from which to compare how networks differ among entities (e.g., subjects, tissue types, or cell types), treatment levels, and times (Fig. 1**B**).

The main merit of a mathematical model is its ability to make a prediction for the future. The qdODEs allow the independent and dependent expression levels of genes to be calculated as long as EI is provided. Thus, for those samples that are not included in our network reconstruction, we can interpolate or extrapolate gene networks based on their EIs. Individualized networks are likely to be associated with clinical and disease phenotypes and, therefore, can be potentially useful for predicting health risk.

#### (v) Biologically meaningful and socially interpretable

Because of bidirectional and signed features, the network can discern distinct patterns of gene interactions (Fig. 1**B**). If two genes facilitate each other by producing factors that promote both parties, then **synergism** occurs. In contrast, an **antagonism** occurs if two genes inhibit each other. **Commensalism** results if one gene promotes its partner but the latter does not affect the former (neutral), while **amensalism** occurs if one gene inhibits the other and the other is neutral. If one gene inhibits the other but the latter promotes the former, then the former exerts **parasitism** to the latter. Conversely, one gene promotes the other but the latter inhibits the former, then the former offers **altruism** to the latter. A lack of any interaction, then, is when two genes coexist and are neutral to each other. These interaction patterns contain the underlying mass, energetic, or signal basis of gene interactions and, therefore, they are more biologically meaningful than the traditional descriptions of genetic epistasis based on statistical tests. A gene may actively manipulate other genes (by promoting or inhibiting the latter) but, meanwhile, may also be passively manipulated by other genes. In networks reconstructed from our approach, one can identify the numbers of such active links and passive links for each gene. If a gene has more active links than passive links, it is regarded as a social gene. If a gene’s active links are more than the average of all genes (i.e., connectivity), then this gene is a core gene that is believed to play a pivotal role in maintaining gene networks. If a gene has less links, including active and passive, than the average, it is a solitary gene.

## Results

### Human vein bypass grafting

Rehfuss et al.^27^ reported a genomic study of infrainguinal vein bypass grafting involving 48 patients, among whom 35 succeeded and 13 failed. To investigate the genomic mechanism underlying graft outcome, transcriptomes of circulating monocytes from patients of success and failure were monitored at pre-operation and at days 1, 7, and 28 post-operation. We selected a subset of genes measured (1,870) that change significantly as a function of time per ANOVA (P < 0.05) for idopGRN reconstruction. Four time points of gene monitoring for 48 patients form 4×48 = 192 samples.

By plotting the expression of individual genes against EI across these samples, we found that each gene’s EI-varying expression is broadly in agreement with the part-whole relationship theory. In Fig. 2, we chose four representative genes for their fitness to the power equation (13). The expression of ADAM9 and LCN2 increases with EI, but the former displays a greater slope of increase (Fig. 2**A**) than does the latter (Fig. 2**B**). In contrast, the expression of PLXNA4 (Fig. 2**C**) and NSUN7 (Fig. 2**D**) decreases with EI, but with different slopes. We used Kim et al.’s functional clustering^47^ to categorize all genes considered into 145 modules each with a distinct EI-varying pattern.

**Figure 2.**
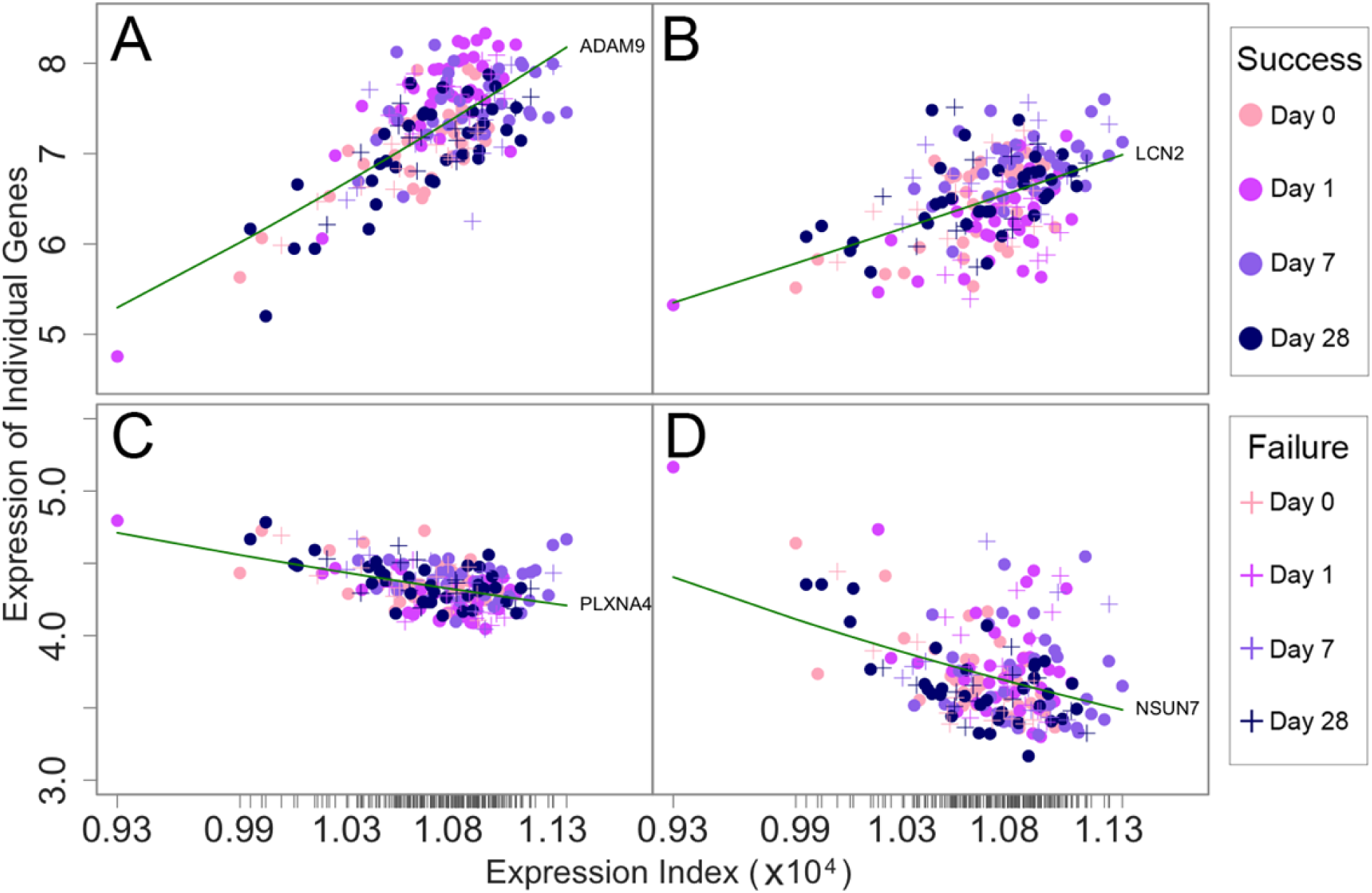
The fitness of a power equation as a function of expression index (EI) (green line) to the observed expression levels of four genes, ADAM9 (**A**), LCN2 (**B**), PLXNA4 (**C**), and NSUN7 (**D**), chosen from the genomic study of human infrainguinal vein bypass grafting, across samples. Samples involve 48 patients, i.e., 35 successes (plus) and 13 failures (circle), multiplied by four time points (including day 0 pre-operation and days 1, 7, and 28 post-operation). Ticks on the x-axis represent the positions of each sample in terms of its EI.

We randomly choose one successful patient (#125) and one failed patient (#205) and compare how they respond to grafting through network alterations. GRNs that specify the alterations of gene co-expression across environmental change are called environment-perturbed GRNs. Figure 3 illustrates graft-perturbed idopGRNs at the module level from pre-operation to days 1 (**A**), 7 (**B**), and 28 (**C**) post-operation, respectively, for #205 (upper panel) and #125 (lower panel). The two patients display some commonalities and differences in terms of their network structure and sparsity. For example, module 53 is a hub that actively regulate many other modules in both success and failure graft-perturbed GRNs. This module only contains an antisense lncRNA gene, C5orf26/EPB41L4A-AS1, located in the 5q22.2 region of the genome [99]. This gene plays a role in the development, activation, and effector functions of immune cells [100]. However, the two networks are remarkably different in many aspects. First, the success network contains more links than the failure network at the early and middle stage of recovery after grafting, but this difference disappears at the late stage of recovery, suggesting that the successful patient can more quickly establish a stable network than the failed patient. Second, the success network from pre-operation to day 1 post-operation is framed by multiple hubs (including not only 53 but also 5, 86, and 109), each displaying strong links with many other modules, but the failure network is only dominated by hub 53 with relatively weak links to other modules. Third, graft-perturbed networks alter more dramatically in topological structure across time for the failed patient than the successful patient.

**Figure 3.**
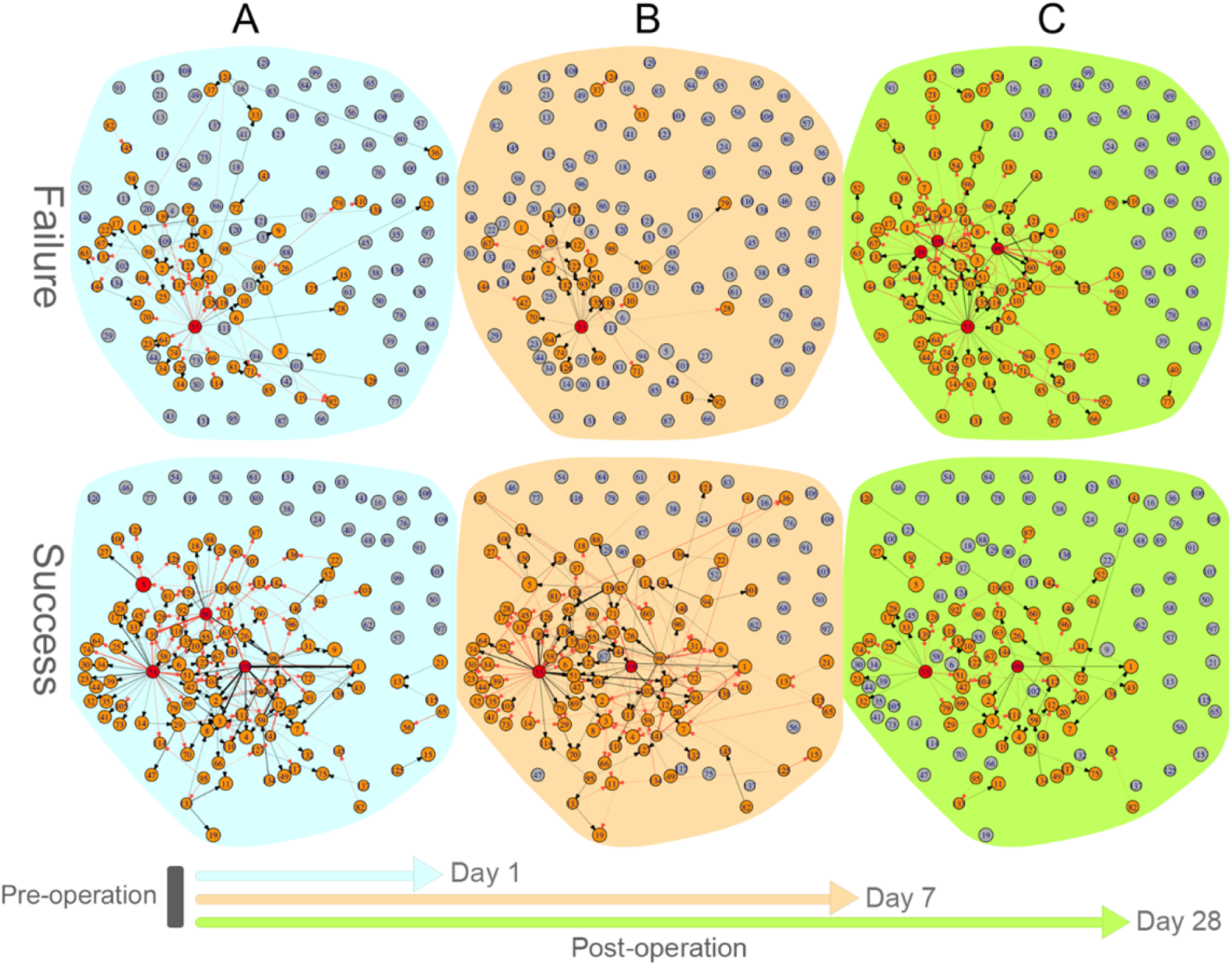
Graft-perturbed networks that code how different gene modules are co-expressed for a failed patient (upper panel) and a successful patient (lower panel) in response to physiological changes from pre-operation to day 1 (**A**), 7 (**B**), and 28 (**C**) post-operation. Numbers in small circles (each denoted as a node of the graph) represent module IDs. Red and blue arrows denote the direction by a gene promotes and inhibits other genes, respectively, and the thickness of an arrowed line is proportional to the strength of promotion or inhibition. A proportion of modules are unlinked, suggesting that they are neutral to each other and other linked genes. Dark red circles denote hub modules with higher connectivity than the average number of links among all modules.

We reconstructed outcome-perturbed networks between successful and failed outcomes at different stages of operation (Fig. 4). We argue that if networks are not associated with graft outcomes, outcome-perturbed networks should be similar structurally preoperatively and post-operation. The outcome-perturbed network prior to operation is dominated primarily by hub module 53, followed by module 124 (Fig. 4**A**), but the outcome-perturbed network at day 1 post-operation involves hubs 53, 124, 109, 59, and 5 (Fig. 4**B**). Module 53 drives the prior network purely through inhibiting other modules, whereas much of its role in the post network is played by promotion. Outcome-perturbed networks at days 7(Fig. 4**C**) and 29 post-operation (Fig. 4D) differ not only from that prior to operation in terms of the number and type of hub modules, but also are sharply contrast to those at day 1 post-operation. Taken together, the genomic mechanisms driving outcome difference can be interrogated by the topology of graft- and outcome-perturbed idopGRNs reconstructed by our approaches.

**Figure 4.**
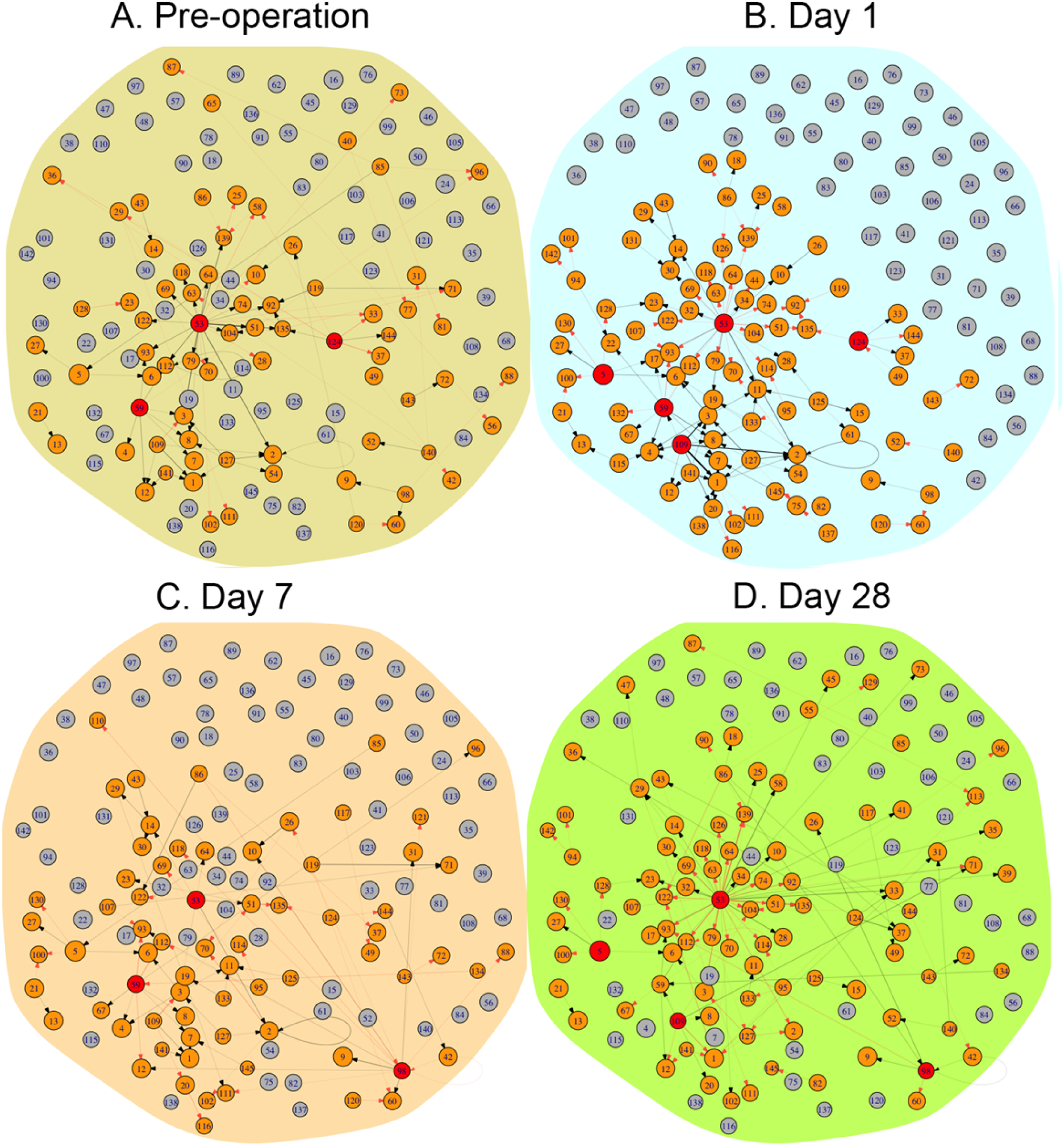
Outcome-perturbed networks that code how different gene modules are co-expressed in response to successful vs. failed patients prior to operation (**A**) and day 1 (**B**), 7 (**C**), and 28 (**D**) post-operation. Numbers in small circles (each denoted as a node of the graph) represent module IDs. Red and blue arrows denote the direction by a gene promotes and inhibits other genes, respectively, and the thickness of an arrowed line is proportional to the strength of promotion or inhibition. A proportion of modules are unlinked, suggesting that they are neutral to each other and other linked genes. Dark red circles denote hub modules with higher connectivity than the average number of links among all modules.

How much a gene is expressed across dynamic networks is determined by its endogenous encoding force and the exogenous influence by other genes. Our approach can dissect the overall expression level of each gene into its independent and dependent expression components. The sign and size of the dependent components can explain how each gene is regulated by other genes in the networks. Four representative modules 20, 27, 118, and 135 exhibit distinct expression patterns across samples, whose underpinnings can be illustrated by drawing the independent and dependent expression curves (Fig. 5). The independent expression of each module increases exponentially with EI, but the slopes of increase vary depending on module type. Modules 20 and 27 are each promoted by other modules, 109, 1, 59 and 115 for the former (Fig. 5**A**) and 5, 53, and 13 for the latter (Fig. 5**B**), both listed in the order of promotion degree. These modules produce accumulative positive dependent effects on the expression of modules 20 and 27, leading the observed expression level of these two focal modules to be higher than their independent expression level across EI gradients. By contrast, the independent expression level of modules 118 and 135 is downshifted by a set of eight modules for the former (Fig. 5**C**) and a set of four modules for the latter (Fig. 5**D**). These two sets of modules inhibit the expression of modules 118 and 135, respectively, producing accumulative negative dependent effects on the focal modules.

**Figure 5.**
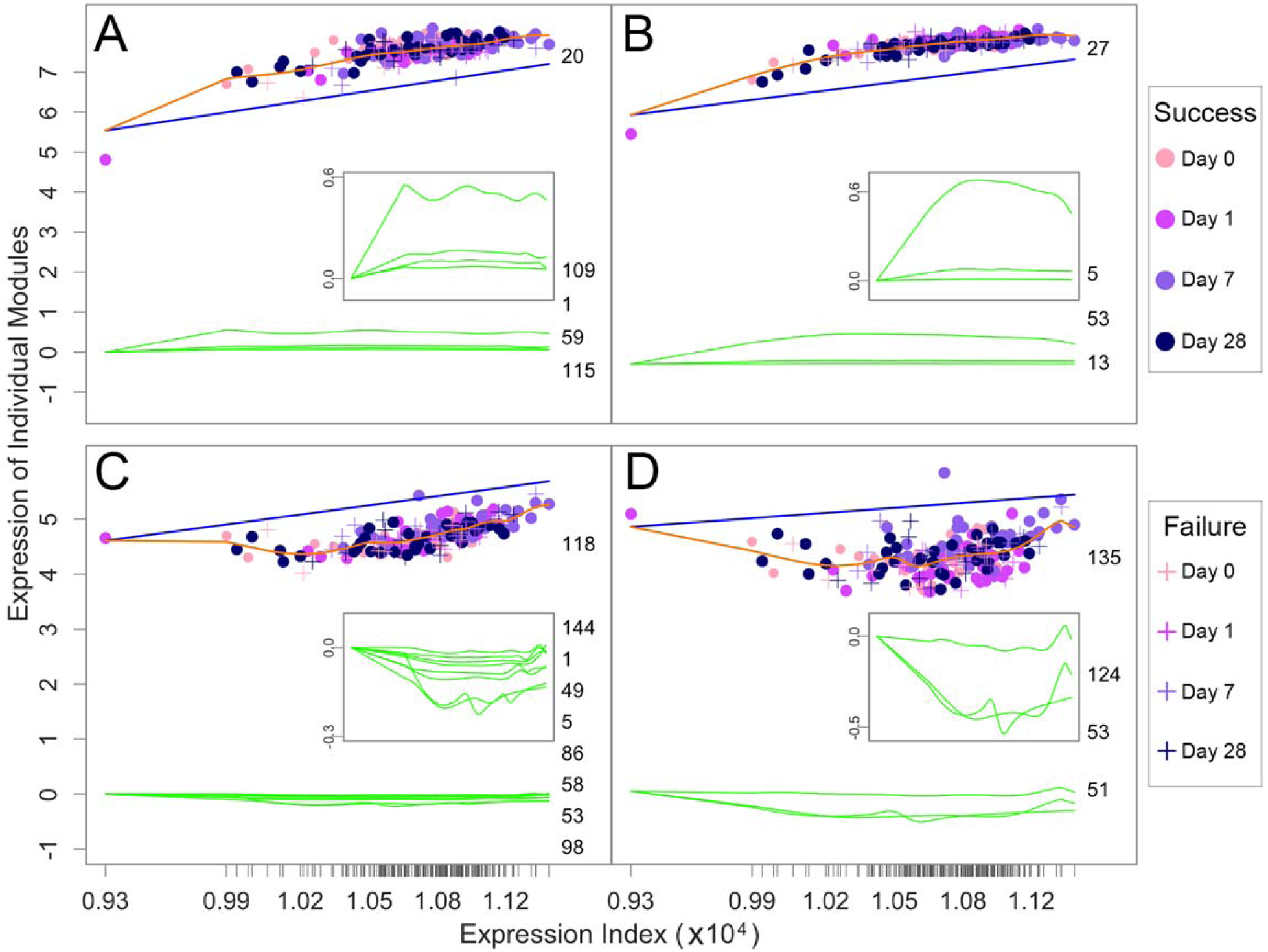
Overall fitted curves of gene expression (orange line) from modules 20 (**A**), 27 (**B**), 118 (**C**), and 135 (**D**) by a system of qdODEs as a function of expression index (EI) in the human vein grafting study. Each dot denotes a sample representing a patient with outcome success (plus) or failure (circle), measured at a time point (day 0 pre-operation and days 1, 7, and 28 post-operation). The overall expression curve of each module is decomposed into its endogenous expression curve (blue line) and exogenous expression curves (green lines) exerted by a set of other modules (listed by their IDs). Exogenous expression curves are better displayed by a small plot within each large plot. Value 0 at y-axis is a cut-off point that describes how a focal module is regulated by other modules: Greater than 0 for promotion, less than 0 for inhibition, and zero for neutrality. Ticks on the x-axis represent the positions of each sample in terms of its EI.

### Rabbit vein bypass graft

We analyzed a second data set of gene expression to validate the usefulness of our approach. The data of microarray genes was collected from a rabbit bilateral vein graft construct^28^. New Zealand white rabbits (weighing 3.0–3.5 kg) of high genetic similarity were treated by bilateral jugular vein interposition grafting and unilateral distal carotid artery branch ligation to create two 6-fold different blood flows. Thousands of genes were monitored on vein grafts, harvested at 2 hours, 1, 3, 7, 14, 30, 90 and 180 days after implantation, under both conditions, high flow and low flow. Each outcome involves three to six rabbits at each time point, which totalize 73 samples. We chose a set of differentially expressed genes (1,395) for idopGRN reconstruction. We calculated the EI of each sample with these genes and plotted the expression of individual genes against EI. EI-varying expression profiles, fitted by a power function (Fig. S1), were clustered into 50 modules (Fig. S1).

We reconstructed module-based idopGRNs of gene co-expression altered from time 2 hours to 1 (A), 30 (B), and 180 days (C) after implantation under high and low flows (Fig. S2). These networks change strikingly in the structure and connectivity across times under both flow conditions. Also, at the same time, idopGRNs differ between high and low flows. Flow-perturbed networks are structurally simple at time 2 hours, but show increasing complexities with time (Fig. S3), suggesting that high and low flows need a time to display their differences. Figure S4 illustrates how the expression of four modules is determined by their endogenous capacity and the exogenous influence of other modules. The overall expression of modules 3 (A), 45 (B), and 38 (D) was observed to be higher than their independent expression because of positive influences exerted by other modules, but module 20 (C) is negatively affected by other modules, making its overall expression lower than independent expression. Taken together, results from the rabbit grafting study support the usefulness of our network inference approach.

### Computer simulation

We will perform computer simulation studies to examine the stability, robustness, and sensitivity of our approach under different scenarios of different sample sizes and measurement errors. We simulated the expression data of *m* genes, *y*_*j*_ *=* (*y*_*j*_(*E*_1_), …, *y*_*j*_(*E*_*N*_)) (*j =* 1, …, *n*), across *N* samples, with *y*_*j*_(*E*_*i*_) varying with *E*_*i*_ (*i =* 1, …, *N*). The EI-varying expression change of gene *j* is specified by an arbitrary form of endogenous expression curve and the sum of arbitrary forms of exogenous curves determined by a set of other genes, plus the residual error of gene *j* in sample *i*, following a multivariate normal distribution with the mean vector **0** and covariance matrix **Σ** whose structure following the AR(1) model. We design different scenarios by changing the number of samples, variance and covariance.

Suppose the expression data of 50 genes across 50, 100, and 200 samples are simulated, respectively. Each gene interacts with a specific set of genes across samples, which are specified by a system of EI-varying qdODEs in Eq. 4. The residual variances and correlation coefficient of gene expression are set as 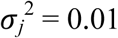 or 0.1 and ρ = 0 or 0.3, respectively. The statistical efficacy of the new approach in terms of gene-gene interaction detection was evaluated by several conventional criteria, including true positive (TP), false positive (FP), true negative (TN), and false negative (FN), from which true positive rates (TPR) and false positive rates (FPR) are calculated by TPR = TP/(TP+FN) and FPR = FP/(FP+TN). In addition, the area under the curve (AUC) of the receiver operating characteristic curve (ROC) was calculated from the coordinates of TPR and FPR. Table S1 gives the results from our simulation studies under different parameter combinations. FPR is very low in every case, suggesting that the approach can be safely used in practice. In general, TPR is reasonably good, but depending on sample size and measurement error. If a small sample size (say 50) is used, we need to improve gene measurement precision to obtain good interaction detection power (say 0.75). If the measurement precision cannot be assured, sample size should be large enough. AUC performs quite well although it also depends on sample size and measurement error.

## Discussion

The past two decades have witnessed countless transcriptional experiments initiated to explore the genomic mechanisms underlying high-order phenotypes for a wide range of organisms. These experiments were designed to monitor gene expression profiles of biological entities under contrast conditions and/or across developmental times. By various comparative analysis and tests, genes expressed differentially under different conditions or over times are identified as biomarkers of phenotypic variation. Cluster analysis was also used to detect distinct patterns of gene expression, facilitating the interpretation of the genomic control over phenotypic or developmental plasticity^28^. However, these widely used standard genomic experiments have not purported to reconstruct gene regulatory networks (GRNs), although these networks play a major role in linking genotype to phenotype^1,2^. The inference of informative GRNs critically relies upon more expensive experiments that are specially designed to produce either perturbed expression data or high-density temporal expression data (Huynh-Thu and Sanguinetti 2018).

In this article, we represent an interdisciplinary approach for reconstructing biologically meaningful GRNs from standard gene expression experiments. How much a gene is expressed in a biological entity is determined by multiple endogenous and exogenous factors. These factors together form the “ecological” component of the entity related to the gene’s overall expression within a network, which can be virtualized as the niche of the gene according to ecology theory^30^. While niche differences maintain the stability of gene networks, the sum of gene-specific niches on an entity reflects the entity’s capacity to supply energy and material for all genes to be expressed. We define the total expression amount of all genes on an entity as the niche index (NI) of the entity. We integrate and contextualize the niche theory of biodiversity (describing how genes are expressed differently across entities) and evolutionary game theory (describing how genes are co-expressed differently across entities) to derive a system of quasi-dynamic ordinary differential equations (qdODEs) with the NI derivative. Such qdODEs specify gene interdependence and interconnection, constructed from any transcriptional experiments involving multiple entities under different treatments, monitored at several key stages and/or across spaces. The optimization solution of these ODEs, through the implementation of variable selection, enables the inference and recovery of informative (encapsulating bidirectional, signed, and weighed links), dynamic (tracing network alterations across spatiotemporal gradients), omnidirectional (capturing all possible links but maintaining the sparsity of networks), and personalized (individualizing networks for each entity) GRNs (idopGRNs).

We incorporate community ecology theory to interpret the biological relevance of idopGRNs. Like the pattern of species-species interaction as a function of resource availability^48^, how one gene interacts with others depends on signal transduction and information flow. The same gene may form a synergistic coexistence with the second gene through cooperation, but may establish an antagonistic relationship with the third gene through competition. The biological underpinnings causing each interaction can be speculated by ecological principles.

We validated the utility of our approach by analyzing gene expression data from surgical patients. Vein bypass grafting is an essential treatment for lower extremity arterial occlusive disease, but only with 30 – 50% success rate^27^. The biological mechanisms underlying the outcome of grafts include cue-induced differentiation of gene expression. We used our approach to reconstruct graft- and outcome-perturbed idopGRNs from 1,870 differentially expressed genes and identified identify key genes and key interactions that cause success vs. failure. As an antisense lncRNA gene, located in the 5q22.2 region of the genome, C5orf26/EPB41L4A-AS1 plays a leadership role in regulating other genes within networks (99). How many genes it regulates, how differently it regulate these genes, and how its regulation responds to grafting and recovery are all potentially important for patients to cure. Based on previous functional studies (100), we postulate that the role of C5orf26/EPB41L4A-AS1 in mediating and activating the gene networks toward cure may be executed through its effects on the development, activation, and effector functions of immune cells. We found more links in the networks of successes than those of failures at the early and middle stage of recovery after grafting. Previous ecological studies show that the number of links, which is usually defined as the complexity of a network^49^, is positively correlated with the stability of the network^50-52^. This thus suggest that the successful patient can more quickly establish a stable network than the failed patient. In conjunction with results from the rabbit vein grafting study, it is suggested that idopGRNs determine grafting outcome by their key genes, structure, complexity, and organization.

Given that complex phenotypes form, develop and alter through genetic networks, computational methods for detecting putative functional relationships between genes are clearly needed. Although extensive efforts have been made to reconstruct various GRNs, most network inference methods cannot provide an omnidirectional and quantitative assessment of network structure and organization. Our approach presented in this article has well resolved these issues, additionally equipping the network reconstruction with biologically meaningful interpretations. Our idopGRNs potentially provide powerful tools to explore various omics data, generate mechanistic hypotheses, and guide further experiments, model development, and analyses. By validating or invalidating various hypotheses experimentally, new scientific discoveries can be made, new insights gained, and new network models revised. Our approach can be refined to accommodate the data features of single cell analysis^53^, which enables idopGRNs to explore an in-depth mechanisms that drive remote biochemical, developmental, and physiological transitions from genotype to phenotype.

## Acknowledgements

This study has been supported by Fundamental Research Funds for the Central Universities (NO. 2015ZCQ-SW-06) and grants U01 HL119178 and NICHD 5R01HD086911-02 from the National Institute of Health.

## Author contributions

C.C. and L. J. designed and implemented the algorithm and performed data analysis and computer simulation. M.W., Y.W., B.C., Z.L. Z.W., and W.H. participated in model derivations, data analysis, and result interpretation. S.B. designed and performed the genomic experiments. R.W. conceived of the idea, supervised the study, and wrote the manuscript with inputs from C.C., L.J., and M.W.

## Online Methods

Here, we describe a statistical procedure for solving a system of qdODEs in Eq 1. By obtaining the maximum likelihood estimates of independent and dependent expression amounts of each gene, idopGRNs can be reconstructed.

### Variable selection for interacting genes

Let **y**_*j*_ = (*y*_*j*_(*E*_1_), …, *y*_*j*_(*E*_*N*_)) denote a vector of observed expression values for gene *j* (*j* = 1, …, *m*) over all samples. The observed expression of gene *j* at sample *i* is expressed as

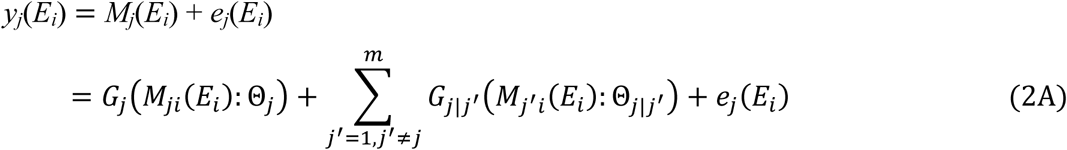

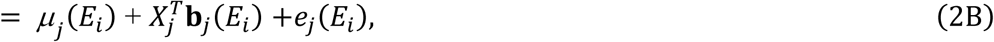

where the overall expression level of focal gene *j, M*_*j*_(*E*_*i*_), includes its independent expression component, *µ*_*j*_(*E*_*i*_) = *G*_*j*_(·) and dependent expression component accumulatively determined by all other genes, 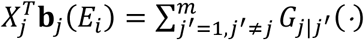; the derivatives of *G*_*j*_(·) and *G*_*j*|*j*′_(·) are *g*_*j*_(·) and *g* _*j*|*j*′_ (·) of ODEs in Eq 1, respectively; and *e*_*j*_(*E*_*i*_) is the measurement error at sample *i*, assumed to be iid with mean zero and variance 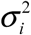 Note that 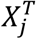 is the vector containing *m* – 1 unities and **b**_*j*_(*E*_*i*_) = (*b*_*j*|1_(*E*_*i*_), …, *b*_*j*|*m*_(*E*_*i*_)) is a vector of the dependent expression of gene *j* determined by all genes, except for gene *j*.

Many nonparametric functions, such as B-spline, regression B-spline, penalized B-spline, local polynomials, or Legendre orthogonal polynomials (LOP), can be used to model independent expression curves, *µ*_*j*_(*E*_*i*_), and dependent expression curves, **b**_*j*_(*E*_*i*_). Chen et al.^12^ have proved statistical properties of B-spline variable selection for solving ODEs. Here, we implement B-spline to fit *µ*_*j*_(*E*_*i*_) and **b**_*j*_(*E*_*i*_) in Eq 2B, allowing orders of nonparametric functions to be gene-dependent and also differ between independent and dependent expression curves. For any gene *j* as a response, there are (*m* – 1) predictors, each of which contributes to the dependent expression of this focal gene through unknown nonparametric dependent parameters **β**_*j*_ **=** (**β**_*j*|1_,…,**β**_*j*|(*j*–1_),**β**_*i*|(*j*+1),_…,**β**_*j*|*m*_). Thus, we have *m* – 1 groups of dependent parameters that reflects the regulation of other genes for the focal gene. We implemented group LASSO^54^ to select those nonzero groups. The group LASSO estimators of dependent parameters, denoted 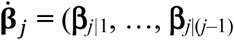, where *d*_*j*_ (≪ *m*) is the number of the most significant genes that interact with gene *j*, can be obtained by minimizing the following penalized weighted least-square criterion,

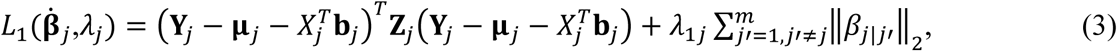

where **y**_*j*_ = (*y*_*j*_(*E*_1_), …, *y*_*j*_(*E*_*N*_)), **y**_*j*_ = (*y*_*j*_(*E*_1_), …, *y*_*j*_(*E*_*N*_)), **µ**_*j*_ = (*µ*_*j*_(*E*_1_), …, *µ*_*j*_(*E*_*N*_)), and **b**_*j*_ = (**b**_*j*_(*E*_1_), …, **b**_*j*_(*E*_*N*_)); *λ*_1*i*_ is a penalty parameter determined by BIC or extended BIC; and **Z**_*j*_ = diag{*z*_*j*_(*E*_1_), …, *z*_*j*_(*E*_*N*_)} where *z*_*j*_(*E*_*i*_) is a prescribed nonnegative weight function on [*E*_1_, *E*_*N*_] with boundary conditions *z*_*j*_(*E*_1_) = *z*_*j*_(*E*_*N*_) = 0. This weight function is used to speed up the rate of convergence.

**Optimizing the topological structure of gene co-expression networks**

Through variable selection, we detect the most significant incoming links (*d*_*j*_ << *m*) for each gene *j* that constitutes the qdODEs of Eq 1. By replacing *m* by *d*_*j*_, these ODEs are modified as

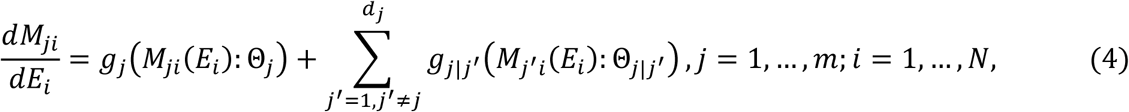

which are a sparse version that represents the full model of incoming links for each gene, but with no constraint on the number of outgoing links and, therefore, the dimension of the network. We formulate a likelihood approach to estimate the modified ODEs. Let **ϕ** = (**μ**;**Σ**) ∈**Φ** denote all model parameters. The likelihood function of given these data is written as

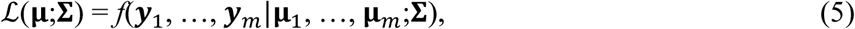

where *f*(·) is the *N*-dimensional *m*-variate normal distribution for *m* gene across *N* samples with mean vector **μ**,

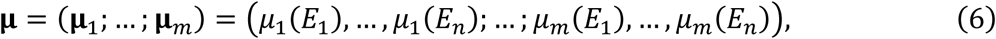

and covariance matrix **Σ**,

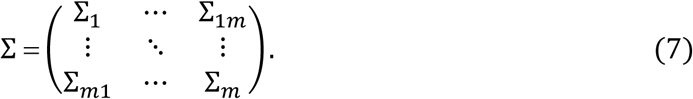

In Eq 6, *µ*_*j*_(*E*_*i*_), the mean value of the expression of gene *j* at sample *i*, whose derivative contains *g*_*j*_(·) and *g*_*j|j*′_(·) specified by the modified qdODEs in Eq 4, is modeled by B-spline function and estimated by standard fourth-order Runge-Kutta algorithms. Since B-spline nonparametric functions are intergrable, we can calculate *G*_*j*_(·) and *G*_*j|j*′_(·). In Eq 7, **Σ**_*j*_ is the sample-dependent covariance matrix of gene *j*, and **Σ** _*jj* ′_ is the sample-dependent covariance matrix between genes *j* and *j*′. We assume that the residual errors of gene expression are independent among samples and that the residual variance of each gene is constant across samples. Thus, **Σ**_*j*_ and **Σ** _*jj* ′_ are structured as 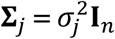 and **Σ** _*jj* ′_ ***=* σ** _*jj*_′ **I**_*n*_ espectively, where 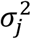 is the residual variance of gene *j* at the same sample, **σ**_*jj* ′_ is the residual covariance of genes *j* and *j*′ at the same sample, and **I**_*n*_ is the identity matrix. However, we implement the first-order autoregressive (AR(1)) model to fit the residual covariances of gene expression among different time points at the same individual55.

All model parameters **ϕ** can obtain their optimal solution by maximizing the likelihood in Eq 5, expressed as

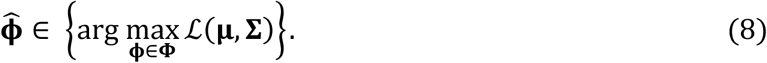

Intuitively, this maximum likelihood optimization implies an optimal topological structure and organization in which genes interact with each other to maximize the expression level of all genes as a whole. This solution of Eq 8 establishes the mathematical formulation of Smith and Price’s evolutionary game theory45.

### Significance test of gene interactions

One important issue for network reconstruction is how to statistically test the significance of edges as the measure of associations between nodes. We propose a likelihood ratio approach for network test. Under the null hypothesis that all microbes are independent from each other, the rate of expression change for each gene can be formulated by a reduced system of ODEs, expressed as

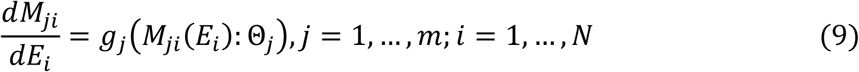

which is contrasted to the full system of ODEs in Eq 4 as the alternative hypothesis stating that at least one gene interaction in the network is significant. We calculate the likelihood values under the null and alternative hypotheses and their log-likelihood ratio (LR) as a test statistic. A network-wise critical threshold can be determined by permutation tests. This procedure includes (i) shuffling sample-varying expression data among genes to make a new data, (ii) calculating the LR value based on this new data, (iii) repeating (i) and (ii) many times (say 1000), and (iv) detecting the 95% percentile of these 1000 LR values which is the cutoff for the significance test of networks.

### Environment-perturbed networks

Genetic networks may be activated when the organism experiences environmental change. Suppose that gene co-expression changes from one sample (say *i*_1_) to next (say *i*_2_) due to differences in the internal environment of samples. The amount of this change can be estimated by integrating the dependent expression component of qdODEs in Eq 4 from 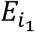 to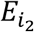, expressed as

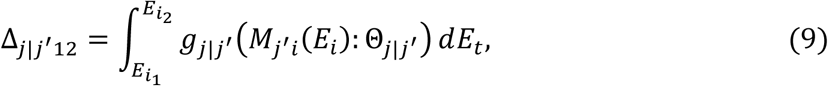

which quantifies the expression difference of gene *j* regulated by gene *j*′ by assuming that sample transport virtually from *i*_1_ to *i*_2_. GRNs reconstructed from _*j/j ′*_ (*j* ≠ *j*′ = 1, …, *m*) reflect the alterations of gene co-expression in response to environmental change, which are called environment-perturbed GRNs. Based on this definition, we can reconstruct treatment-, outcome-, development, or signal-perturbed networks to better understand the genomic mechanisms underlying cellular, physiological, and ecological processes.

## Supplementary Figure Legends

**Figure S1.** The fitness of a power equation as a function of expression index (EI) (green line) to the observed expression levels of four genes, BC011754 (**A**), AB007963 (**B**), BC016908 (**C**), and NM_005103 (**D**) across 73 rabbit samples. Samples include three to six rabbits under each of two blood flows, low (purple circles) and high (dark circles), measured at each of eight time points (hour 2 and days 1, 3, 7, 14, 30, 90, and 180) post-operation. Ticks on the x-axis represent the positions of each sample in terms of its EI.

**Figure S2.** Development-perturbed networks at the module level under low flow (upper panel) and high flow (lower panel) of rabbit vein grafting experiment in response to developmental stimuli from hour 2 to day 1 (**A**), day 30 (**B**), and 180 (**C**) post-operation. Numbers in small circles (each denoted as a node of the graph) represent module IDs. Red and black arrows denote the direction by a gene promotes and inhibits other genes, respectively, and the thickness of an arrowed line is proportional to the strength of promotion or inhibition. A proportion of modules are unlinked, suggesting that they are neutral to each other and other linked genes. Dark red circles denote hub modules with higher connectivity than the average number of links among all modules.

**Figure S3.** Flow-perturbed networks at the module level from slow to high flows of grafted rabbits at hour 2 (**A**), day 1 (**B**), day 30 (**C**), and day 180 (**D**) post-operation. Numbers in small circles (each denoted as a node of the graph) represent module IDs. Red and black arrows denote the direction by a gene promotes and inhibits other genes, respectively, and the thickness of an arrowed line is proportional to the strength of promotion or inhibition. A proportion of modules are unlinked, suggesting that they are neutral to each other and other linked genes. Dark red circles denote hub modules with higher connectivity than the average number of links among all modules.

**Figure S4.** Overall fitted curves of gene expression (orange line) from modules 3 (**A**), 20 (**B**), 45 (**C**), and 48 (**D**) by a system of qdODEs as a function of expression index (EI) in the rabbit vein grafting experiment. Each dot denotes a sample representing a rabbit under a blood flow, low (purple) or high (dark), measured at a time point (hour 2 and days 1, 3, 7, 14, 30, 90, and 180) post-operation. The overall expression curve of each module is decomposed into its endogenous expression curve (blue line) and exogenous expression curves (green lines) exerted by a set of other modules (listed by IDs). Exogenous expression curves are better displayed by a small plot within each large plot. Value 0 at y-axis is a cut-off point that describes how a focal module is regulated by other modules: Greater than 0 for promotion, less than 0 for inhibition, and zero for neutrality. Ticks on the x-axis represent the positions of each sample in terms of its EI.

**Table S1.**
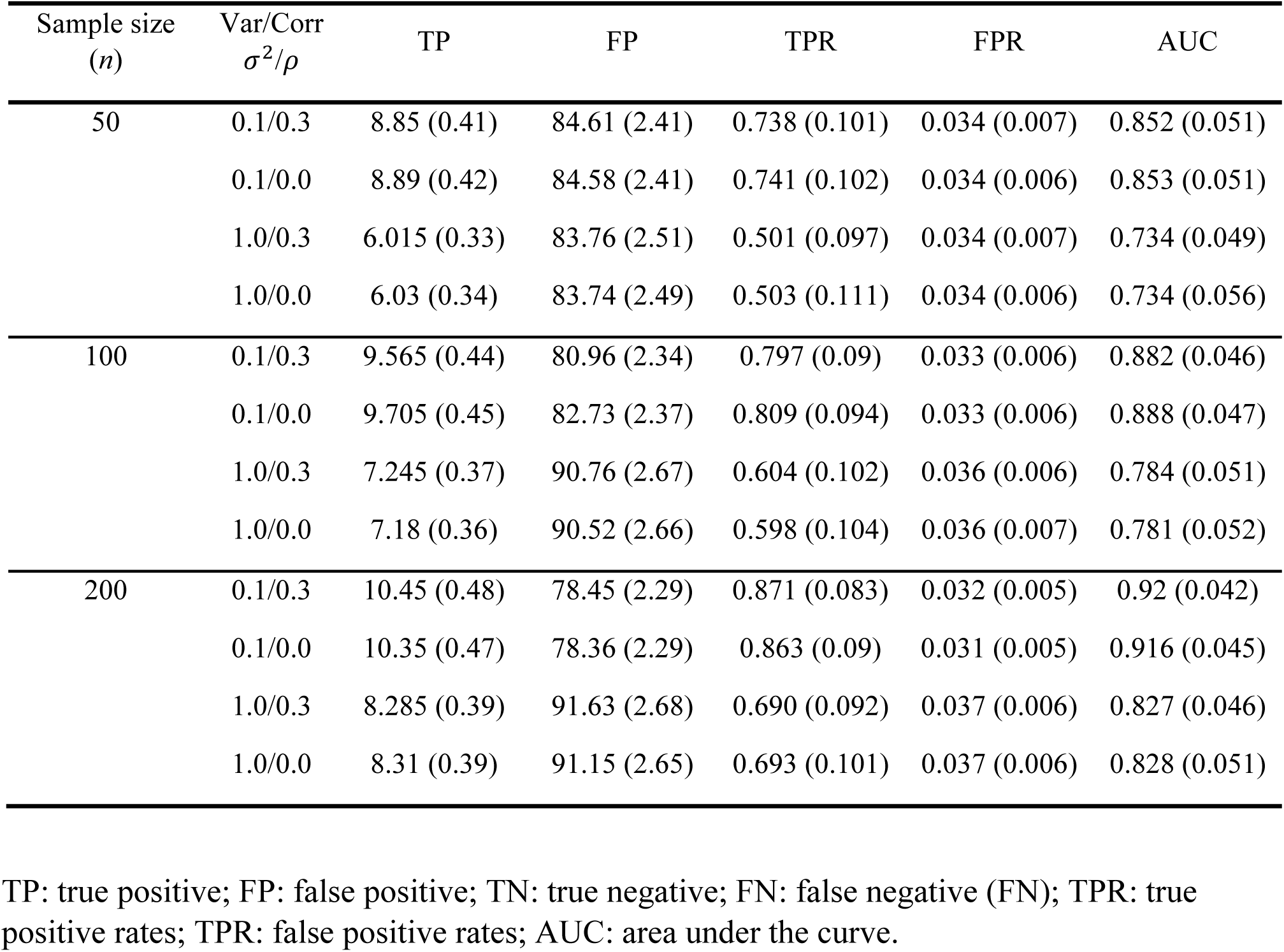
Statistical properties of idopGRN reconstruction under different simulation scenarios. Numbers in parentheses are the standard deviations.

## References

1. Gardner TS, di Bernardo D, Lorenz D, Collins JJ. Inferring genetic networks and identifying compound mode of action via expression profiling. Science 2003;301:102–105.

2. Costanzo M, VanderSluis B, Koch EN, Baryshnikova A, Pons C, Tan G, Wang W, Usaj M, Hanchard J, Lee SD et al. A global genetic interaction network maps a wiring diagram of cellular function. Science 2016;353:pii: aaf1420.

3. Karlebach G, Shamir R. Modelling and analysis of gene regulatory networks. Nat Rev Mol Cell Biol 2008;9:770–780.

4. Oates CJ, Amos R, Spencer SEF. Quantifying the multi-scale performance of network inference algorithms. Stat Appl Genet Mol Biol 2014;13:611–631.

5. Chan TE, Stumpf MPH, Babtie AC. Gene regulatory network inference from single-cell data using multivariate information measures. Cell Syst 2017;5(3):251–267.

6. Bornholdt S. Boolean network models of cellular regulation: prospects and limitations. J Roy Soc Interf 2008;5:S85–S94.

7. Werhli AV, Husmeier D. Reconstructing gene regulatory networks with Bayesian networks by combining expression data with multiple sources of prior knowledge. Stat Appl Genet Mol Biol 2007;6:1–47.

8. Stuart JM, Segal E, Koller D, Kim SK. A gene-coexpression network for global discovery of conserved genetic modules. Science 2003;302:249–255.

9. Wang J, Chen B, Wang Y, Wang N, Garbey M, Tran-Son-Tay R, Berceli SA, Wu RL. Reconstructing regulatory networks from the dynamic plasticity of gene expression by mutual information. Nucleic Acids Res 2013;41:e97.

10. Han SW, Chen G, Cheon M-S, Zhong H. Estimation of directed acyclic graphs through two-stage adaptive Lasso for gene network inference. J Am Stat Assoc 2016;111:1004–1019.

11. Proulx SR, Promislow DE, Phillips PC. Network thinking in ecology and evolution. Trends Ecol Evol 2005;20:345–353.

12. Ghahramani, Z. Learning Dynamic Bayesian Networks. Lecture Notes in Computer Science 1998;1387, 168–197.

13. Perrin BE, Ralaivola L, Mazurie A, Bottani S, Mallet J, d’Alché-Buc F. Gene networks inference using dynamic Bayesian networks. Bioinformatics 2003; Suppl 2:ii138–48.

14. Zou M, Conzen SD. A new dynamic Bayesian network (DBN) approach for identifying gene regulatory networks from time course microarray data. Bioinformatics 2005;21:71–79.

15. Lu T, Liang H, Li H, Wu H. High-dimensional ODEs coupled with mixed-effects modeling techniques for dynamic gene regulatory network identification. J Am Stat Assoc 2011;106:1242–1258.

16. Wu H, Lu T, Xue H, Liang H. Sparse additive ordinary differential equations for dynamic gene regulatory network modeling. J Am Stat Assoc 2014;109:700–716.

17. Henderson J, Michailidis G. Network reconstruction using nonparametric additive ODE models. PLoS ONE 2014;9(4):e94003.

18. Chen S, Shojaie A, Witten D. Network reconstruction from high-dimensional ordinary differential equations. J Am Stat Assoc 2017;112:1697–1707.

19. Swain PS, Elowitz MB, Siggia ED. Intrinsic and extrinsic contributions to stochasticity in gene expression. Proc Natl Acad Sci U S A 2002;99(20):12795–12800.

20. Raj A, van Oudenaarden A. Nature, nurture, or chance: stochastic gene expression and its consequences. Cell 2008;135(2):216–226.

21. Grundberg E, Small KS, Hedman ÅK, Nica AC, Buil A, Keildson S, Bell JT, Yang TP, Meduri E, et al. Mapping cis- and trans-regulatory effects across multiple tissues in twins. Nat Genet 2012;44:1084–1089.

22. Angulo MT, Moreno JA, Lippner G, Barabási AL, Liu YY. Fundamental limitations of network reconstruction from temporal data. J R Soc Interf 2017;14(127).

23. Huynh-Thu V, Sanguinetti G. Gene regulatory network inference: an introductory survey arXiv preprint (2018).

24. Levine JM, HilleRisLambers J. The importance of niches for the maintenance of species diversity. Nature 2009;461:254–257.

25. Tan JQ, Kelly CK, Jiang L. Temporal niche promotes biodiversity during adaptive radiation. Nat Commun 2013;4:2102.

26. Zuppinger-Dingley D, Schmid B, Petermann J, Yadav V, De Deyn GB, Flynn D, Dan FB. Selection for niche differentiation in plant communities increases biodiversity effects. Nature 2014; 515:108–111.

27. Rehfuss JP, DeSart KM, Rozowsky JM, O’Malley KA, Moldawer LL, Baker HV, Wang YQ, Wu RL, Nelson PR, Berceli SA. Hyperacute monocyte gene response patterns are associated with lower extremity vein bypass graft failure. Circ Genom Precis Med 2018;11(3): e001970.

28. Wang YQ, Xu M, Wang Z, Tao M, Wang L, Zhu J, Li RZ, Berceli SA, Wu RL. How to cluster gene expression dynamics in response to environmental signals. Brief Bioinform 2012;13:162–174.

29. Elton CS. Animal Ecology. London: Sidwich & Jackson (1927).

30. Pocheville A. The Ecological Niche: History and Recent Controversies. In Heams, Thomas; Huneman, Philippe; Lecointre, Guillaume; et al. Handbook of Evolutionary Thinking in the Sciences. Dordrecht: Springer. pp. 547–586 (2015).

31. Hui C. Carrying capacity, population equilibrium, and environment’s maximal load. Ecol Model 2006;192:317–320.

32. Finlay KW, Wilkinson GN. The analysis of adaptation in a plant breeding program. Aust J Agr Res 1963;14:742–754.

33. Lobell DB, Roberts MJ, Schlenker W, Braum N, Little BB, Rejesus RM, Hammer GL. Greater sensitivity to drought accompanies maize yield increase in the U.S. Midwest. Science 2014;344:516–519.

34. Pereira FC, Berry D. Microbial nutrient niches in the gut. Environ Microbiol 2017;19(4): 1366–1378.

35. Peterson AT, Soberôn J, Pearson RG, Anderson RP, Martínez-Meyer E, Nakamura M, Araújo MP. Species-environment relationships. In: Ecological Niches and Geographic Distributions (MPB-49). Princeton University Press. p. 82 (2011).

36. McConnaughay KDM, Coleman JS. Biomass allocation in plants: Ontogeny or optimality? A test along three resource gradients. Ecology 1999;80:2581–2593.

37. Xu S, Li Y, Wang G. Scaling relationships between leaf mass and total plant mass across Chinese forests. PLoS ONE 2014;9(4):e95938.

38. Gayon J. History of the concept of allometry. Amer Zool 2000;40:748–758.

39. Shingleton A. Allometry: The study of biological scaling. Nat Ed Knowl 2000;3(10):2.

40. McFarland DJ. Decision-making in animals. Nature 1977;269:15–21.

41. Dias-Ferreira E, Sousa JC, Melo I, Morgado P, Mesquita AR, Cerqueira JJ, Costa RM, Sousa N. Chronic stress causes frontostriatal reorganization and affects decision-making. Science 2009;325:621–625.

42. Park H, Lee D, Chey J. Stress enhances model-free reinforcement learning only after negative outcome. PLoS ONE 2017;12(7):e0180588.

43. Friedman A, Homma D, Bloem B, Gibb LG, Amemori KI, Hu D, Delcasso S, Truong TF, Yang J, Hood AS et al. Chronic stress alters striosome-circuit dynamics, leading to aberrant decision-making. Cell 2017;171:1191–1205.

44. von Neumann J, and Morgenstern S. Theory of Games and Economic Behavior. Princeton University Press, Princeton (1944).

45. Smith JM, Price GR. The logic of animal conflict. Nature 1973;246:15–18.

46. Hofbauer J, Sigmund K. Evolutionary Games and Population Dynamics. Cambridge University Press, Cambridge, UK (1998).

47. Kim B-R, Zhang L, Berg A, Fan J, Wu RL. A computational approach to the functional clustering of periodic gene expression profiles. Genetics 2008;180:821–834.

48. Vellend M. 2010. Conceptual synthesis in community ecology. Q Rev Biol 85: 183–206.

49. MacArthur R. Fluctuations of animal populations and a measure of community stability. Ecology 1955;36:533–536.

50. Arnold JS. Constraints on phenotypic evolution. Am Nat 1992;140:S85–S107.

51. Debat V, David P. Mapping phenotypes: canalization, plasticity and developmental stability. Trends Ecol Evol 2001;16:555–561.

52. Wagner A. Robustness and Evolvability in Living Systems. Princeton University Press (2005).

53. Luijk R, Dekkers KF, van Iterson M, Arindrarto W, Claringbould A, Hop P et al. Genome-wide identification of directed gene networks using large-scale population genomics data. Nat Commun 1018;9(1):3097.

## References

54. Yuan M, Lin Y. Model selection and estimation in regression with grouped variables. J Roy Stat Soc Ser B 2006;68:49–67.

55. Zhao W, Hou W, Littell RC, Wu RL. Structured antedependence models for functional mapping of multivariate longitudinal quantitative traits. Stat Appl Genet Mol Biol 2005;4:Issue 1.

